# pH-Controlled chemoselective rapid azo-coupling reaction (CRACR) enables global profiling of serotonylation proteome in cancer cells

**DOI:** 10.1101/2024.05.10.593574

**Authors:** Nan Zhang, Jinghua Wu, Shuaixin Gao, Haidong Peng, Huapeng Li, Connor Gibson, Sophia Wu, Jiangjiang Zhu, Qingfei Zheng

## Abstract

Serotonylation has been identified as a novel protein post-translational modification for decades, where an isopeptide bond is formed between the glutamine residue and serotonin through transamination. Transglutaminase 2 (also known as TGM2 or TGase2) was proven to act as the main “writer” enzyme for this PTM and a number of key regulatory proteins (including small GTPases, fibronectin, fibrinogen, serotonin transporter, and histone H3) have been characterized as the substrates of serotonylation. However, due to the lack of pan-specific antibodies for serotonylated glutamine, the precise enrichment and proteomic profiling of serotonylation still remain challenging. In our previous research, we developed an aryldiazonium probe to specifically label protein serotonylation in a bioorthogonal manner, which depended on a pH-controlled chemoselective rapid azo-coupling reaction (CRACR). Here, we report the application of a photoactive aryldiazonium-biotin probe for the global profiling of serotonylation proteome in cancer cells. Thus, over 1,000 serotonylated proteins were identified from HCT 116 cells, many of which are highly related to carcinogenesis. Moreover, a number of modification sites of these serotonylated proteins were determined, attributed to the successful application of our chemical proteomic approach. Overall, these findings provided new insights into the significant association between cellular protein serotonylation and cancer development, further suggesting that to target TGM2-mediated monoaminylation may serve as a promising strategy for cancer therapeutics.

**TOC:** 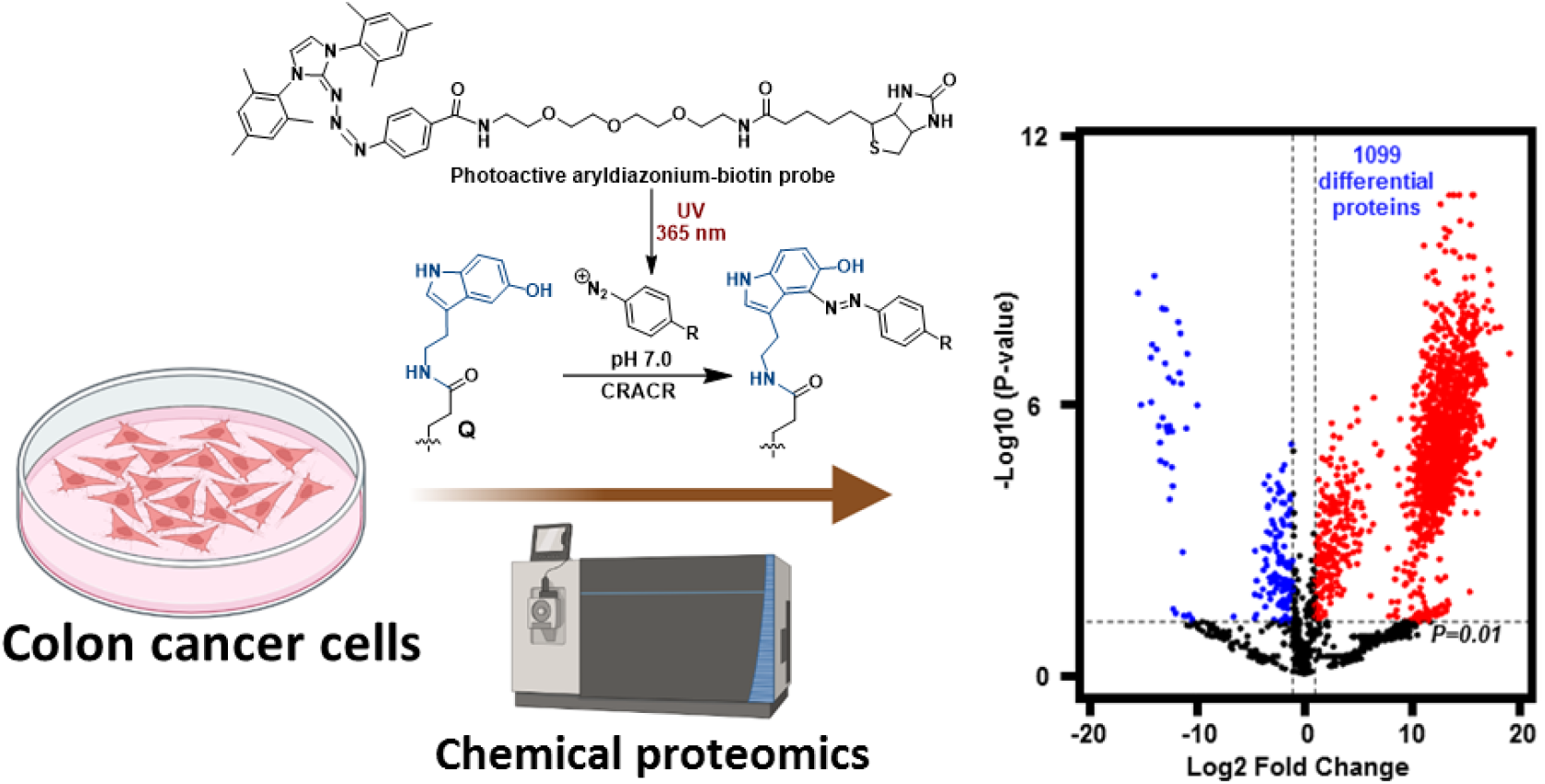

## Introduction

The discovery of serotonylation on protein glutamine (Q) residues dates back to the late 1950s.^1^ In this unique yet ubiquitous post-translational modification (PTM), an isopeptide bound is formed between the primary amine of serotonin [also referred to as 5-hydroxytryptamine (5-HT)] and side-chain amide of proteinogenic glutamine through transamination catalyzed by transglutaminases.^2^ Transglutaminase 2 (also known as TGM2 or TGase2) has been shown to be the major “writer” enzyme for this PTM process, due to its ubiquitous expression in diverse types of organs and cells.^3-5^ A variety of key regulatory proteins (*e*.*g*., small GTPases,^6^ fibronectin,^7^ fibrinogen,^8^ serotonin transporter,^9^ histone H3,^10^ *etc*) have been identified to undergo serotonylation, which plays a role in signaling transduction and gene transcription. One well-established example is the emerging epigenetic mark, H3Q5 serotonylation, which has a significant impact on transcription regulation via multiple mechanisms.^11-14^

Even though a number of proteins have been individually identified as serotonylation substrates, the global profiling of cellular serotonylated proteome is still challenging. Due to the relatively low abundance of serotonylation, suitable tools (such as pan-specific antibodies) are needed for the specific enrichment of serotonylated proteins before mass spectrometry (MS)-based proteomics analysis.^15^ However, till now, there is no report regarding the successful development of pan antibodies against serotonylated glutamine. Therefore, to enrich the cellular serotonylated proteome, an alkyne tag-containing probe, 5-propargyltryptamide (5-PT),^16^ was utilized as a serotonin mimic to track and visualize protein serotonylation,^10,16,17^ while this strategy has obvious limitations. In this copper-catalysed azide-alkyne cycloaddition (CuAAC)-based chemical biology approach, 5-PT needs to be exogenously added to the cultured cells, which cannot be used for *in situ* labeling of endogenous serotonylation within tissue samples. Moreover, blocking the key functional C5-hydroxyl group of serotonin by alkyne also significantly influence its transport across the plasma membrane and signaling transduction. Therefore, in a reported proteomic profiling of protein serotonylation, excess 5-PT and exogenous recombinant TGM2 enzyme were added to the cell lysates.^18^ This strategy may not be ideal for enriching and investigating the endogenously serotonylated proteome, and thus only 46 proteins were identified undergoing serotonylation from SW480 cells.^18^ More advanced chemical proteomics approaches are needed to fulfill the global profiling of cellular serotonylated proteome.

Recently, we developed a photoactive aryldiazonium-biotin probe to label and enrich serotonylated proteins in a bioorthogonal manner,^14^ which relied on a pH-controlled chemoselective rapid azo-coupling reaction (CRACR).^19-21^ This pH-controlled bioorthogonal reaction has been applied to label 5-hydroxytryptophan (5-HTP) and tyrosine under different pH values. Specifically, when the pH is over 9, CRACR occurs on tyrosine residues, while the C4 position of 5-hydroxyindole moieties (such as serotonin and 5-HTP) is rapidly substituted by aryldiazonium under pH 6-7 (**Fig. S1**), due to the electron-donating effect.^19-21^ Utilizing this chemical biology tool, we discovered for the first time that histone H3 serotonylation was accumulated in colorectal cancer cells and regulated gene transcription through multiple mechanisms.^14^ Here, we applied our photoactive aryldiazonium-biotin probe and took advantage of the pH-controlled CRACR for the global profiling of serotonylation proteome in colon cancer cells (**Fig. 1**). Over 1,000 serotonylated proteins were thus enriched and identified, many of which are closely related to cancer development. Kyoto Encyclopedia of Genes and Genomes (KEGG) pathway and interaction network analyses further showed that these serotonylated proteins were involved in diverse essential and fundamental biological processes (such as DNA repair, chromatin remodeling, hormone biosynthesis, mRNA metabolism, transcription, *etc*). Notably, there are a number of overlapping proteins between the serotonylation and dopaminylation proteome,^22^ which further confirms the activation-substitution catalytic mechanism of TGM2 and the microenvironment-driven nature of monoaminylation.^17^ Finally, we identified a variety of modification sites on the key serotonylated proteins, due to the successful application of our aryldiazonium-biotin probe. Overall, the CRACR-based chemical proteomic profiling of serotonylated proteins in this study shed light on the ubiquitous existence of serotonylation in cancer cells and the potential regulatory roles it plays in cellular signaling transduction.

**Figure 1.**
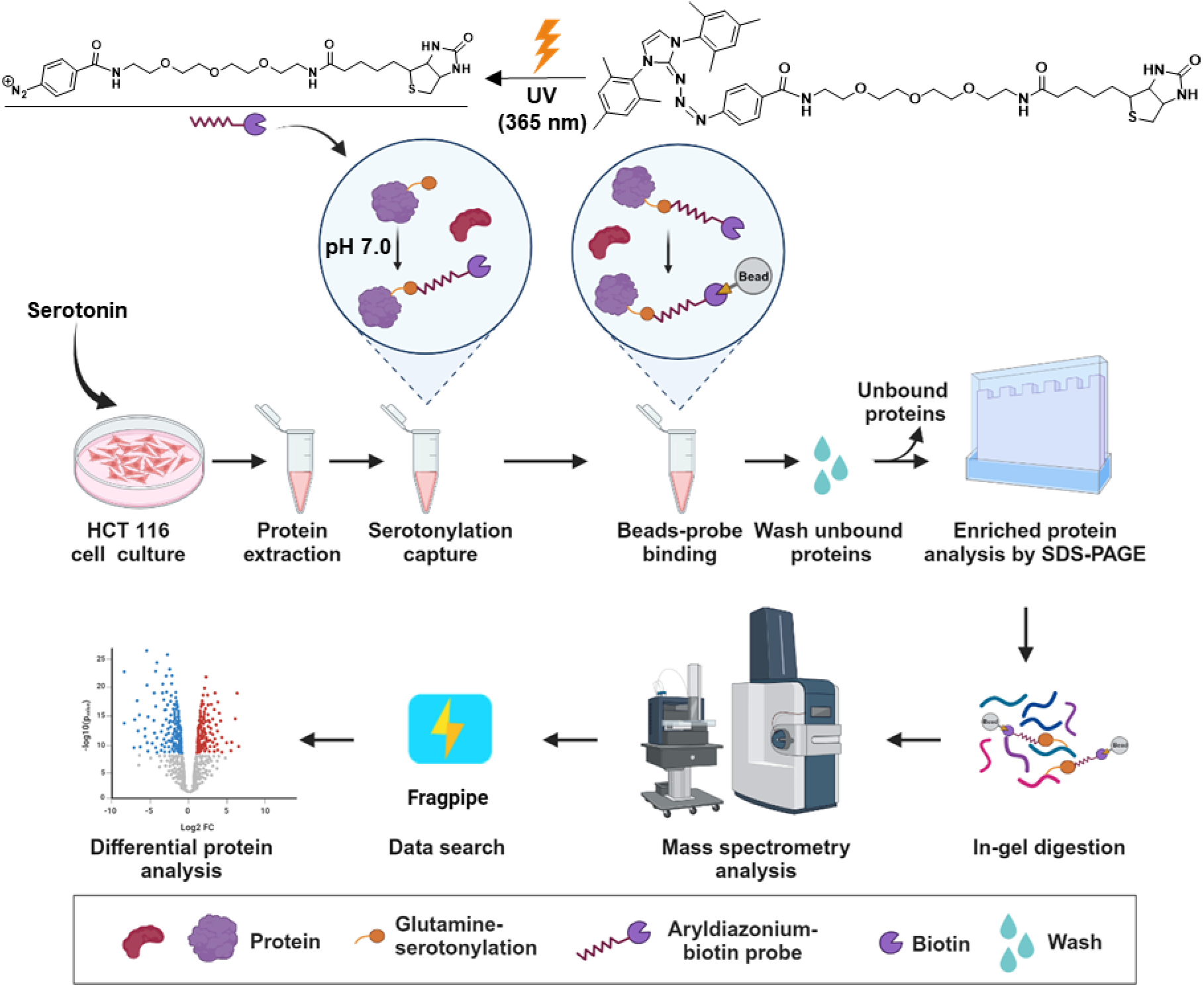
Workflow of the chemical proteomics approach developed in this study for the global profiling of protein serotonylation in colon cancer cells and the structure of aryldiazonium-biotin probe used in this work.

## Experimental section

### Colorectal cancer cell culture

The colorectal cancer cell lines were cultured based on the previously reported protocol with slight modifications.^22^ Briefly, HCT 116 cells were cultured in Dulbecco’s Modified Eagle Medium (DMEM) supplemented with 10% fetal bovine serum (FBS) and 1% sodium pyruvate under 5% CO_2_ at 37 °C. Upon reaching 80% confluence, 10 mL of medium containing 50 µL of 100 mM serotonin hydrochloride was added to achieve a final concentration of 0.5 mM. The medium was gently mixed, and cells were collected through centrifugation after 12 hours of culture. The cell pellets were washed three times using 1X Dulbecco’s Phosphate-Buffered Saline (DPBS) buffer before being further processed.

### Probe labeling and enrichment of serotonylated proteins

The cytoplasmic and nuclear proteins of HCT 116 cells were extracted using the Cytoplasmic and Nuclear Protein Extraction Kit (BOSTER) in accordance with the provided instructions. Subsequently, 140 µL of the protein samples were separately distributed into two 1.5mL tubes, designated as the experimental and control groups, for the further probe labeling reactions. To initiate the labeling reaction, the aryldiazonium-biotin probe was firstly dissolved in dimethyl sulfoxide (DMSO) and activated by ultraviolet (UV) at a wavelength of 365 nm as described before.^14^ Briefly, the probe solution was photolyzed for 1 min using a 43 W UV light (Kessil PR160L-370nm). Thereafter, 85 µL of the extracted protein samples were incubated with 10 µL of 1 M phosphate buffer (pH 7.0). Subsequently, 5 µL of 20 mM UV-activated probe solution was added to the mixture with a final volume of 100 µL, followed by a 20-minute incubation on ice. Unreacted diazonium was quenched by adding 1 mM 5-HTP. In the meantime, a control group was prepared by adding 5 µL of 20 mM biotin solution (instead of the probe), which had been pre-treated with UV.

The samples were then desalted using Zeba spin columns (Thermo Fisher Scientific) to get rid of the small molecules from the proteins. The streptavidin magnetic beads (Thermo Fisher Scientific) employed for enrichment underwent an initial blocking step with 1% bovine serum albumin (BSA) dissolved in 1X phosphate buffer (pH 8.0) on a rotator for 1 hour at room temperature. Subsequently, the beads were washed three times with 1X phosphate buffer for 10 minutes each. The labeled protein samples were then incubated with 40 μL streptavidin beads in 1X phosphate buffer on a rotator at 4 °C overnight to facilitate the enrichment of BCN probe-labeled proteins. Following the incubation, the beads were collected using a magnetic stand and subjected to a series of washes. Initially, the beads were washed with 1 mL of 1% NP40, 0.1% SDS, and 4 M urea, where each rotation lasted for 1 hour at room temperature. Subsequently, the beads were washed with 1X PBST (PBS containing 0.1% Tween 20 detergent) for 10 minutes to remove urea.

### SDS-PAGE analysis and silver staining

These steps were performed according to our previous protocols.^22^ Briefly, the processed streptavidin beads were mixed with 20 µL of 2X loading buffer (Thermo Fisher Scientific) and heated at 98°C for 10 minutes. The samples were then centrifuged at 16000X g for 10 minutes, and the resulting supernatant was loaded onto the SDS-gel for SDS-PAGE analysis. For visualization, silver staining was performed using the Pierce™ Silver Stain for Mass Spectrometry kit, adhering to the provided instructions.

### In-gel digestion

This step was performed according to our previous protocols.^22^ Briefly, the protein lanes on the gel corresponding to the experimental and control samples were excised by using a surgical blade. Each gel strip was then placed into a 1.5 mL centrifuge tube and sectioned into small fragments (0.5 × 0.5 mm). The decolorizing liquid (buffer A and B in the silver staining kit) was added to immerse the gel particles until the yellow color faded, and the resulting supernatant was discarded. Following this, the gel fragments were treated with 300 μL of 50 mM NH_4_HCO_3_ for 10 minutes. After removing the liquid, 300 µL of 100% acetonitrile was added, and the tubes were placed on a shaker for 15 minutes to dehydrate the gel fragments. The supernatant was then removed, and the gel was air-dried. Thereafter, the gel fragments were incubated with a solution containing 100 mM NH_4_HCO_3_ and 10 mM tris(2-carboxyethyl)phosphine (TCEP) for 30 minutes at room temperature. The supernatant was removed, and the gel fragments were dehydrated again using 300 uL of 100% acetonitrile. Further treatment involved adding 100 μL of a solution containing 55 mM iodoacetamide in 100 mM NH_4_HCO_3_, followed by an alkylation reaction in the dark for 30 minutes. After removing the supernatant, each tube was treated with 300 μL of 100 mM NH_4_HCO_3_ at room temperature for 15 minutes on a shaker. Before enzyme lysis, another dehydration step was conducted with 300 μL of 100% acetonitrile. Each tube was then filled with 200 μL (sufficient to cover the gel particles) of a 50 mM NH_4_HCO_3_ solution containing 2 μg of trypsin (Fisher Scientific) and incubated in a temperature-controlled shaker at 37 °C for approximately 20 hours. Following enzymatic digestion, the liquid was transferred to another tube, and 150 μL of a prepared buffer (containing 5% formic acid and 50% acetonitrile) was added for peptide extraction for 20 minutes at room temperature. The tubes were shaken for 1 minute, and this extraction step was repeated three times. The final extraction was conducted using 100% acetonitrile. All the liquid was collected into a 1.5 mL centrifuge tube, followed by vacuum drying. The dried powder resulting from the previous steps was reconstituted by adding 600 µL of 0.1% formic acid aqueous solution. The follow-up desalting was carried out using the peptide desalting spin columns (Pierce), following the product instructions. Finally, the desalted peptides were vacuum-dried in preparation for further LC-MS analyses.

### Mass Spectrometric Analysis

The peptide samples were dissolved in 0.1% formic acid and then analyzed by using two types of mass spectrometers: Orbitrap Fusion Tribrid Mass Spectrometer (Thermo Fisher) for the analysis of protein differentiation; Hybrid Trapped Ion Mobility Spectrometry (TIMS) Quadrupole Time-of-flight Mass Spectrometer (Bruker timsTOF Pro) for the serotonylation site analysis.

For the analysis using Orbitrap Fusion, a CaptiveSpray nanoelectrospray ion source and a nanoElute nanoflow LC system (Bruker) were employed. Peptides were separated on an Acclaim™ PepMap™ 100 C18 HPLC Column (Thermo, 75µm×25cm, 2.0µm C18) using a linear acetonitrile gradient (0%-2% over 5 minutes, 2%-16% over 100 minutes, 16%-25% over 20 minutes, 25%-85% over 1 minute, held at 85% for 4 minutes, 85%-2% over 1 minute, and finally held at 2% for 14 minutes) at a constant flow of 300 nL/min. Peptides were detected using a data-dependent HCD method.

For the analysis using timsTOF Pro, which was equipped with Ion Mobility FAIMS Pro seamless connection and an UltiMate 3000 HPLC, samples were loaded onto a C18 column (Ionopticks, 75 µm X 250 mm). Peptide separation was achieved at a constant flow of 300 nL/min with a linear acetonitrile gradient, starting at 2% and increasing to 22% over 94 minutes. The gradient was then adjusted to 37% acetonitrile for 10 minutes, followed by 100% acetonitrile for 6 minutes. A final step maintained 100% acetonitrile for an additional 10 minutes to ensure complete peptide analysis. Peptides were detected using a data-dependent CID method in a PASEF (Parallel Accumulation and Serial Fragmentation) mode, with 10 PASEF scans performed during each topN acquisition cycle.

### Biochemical validation of serotonylation sites

To validate the serotonylation sites identified by using mass spectrometry, site-directed mutagenesis, immunoprecipitation (IP), and immunoblotting were applied. Firstly, the EGFP-tagged RRP12Q699A mutation was cloned by site-directed mutagenesis using pT7-EGFP-C1-HsRRP12_K as the template, which was a gift from Elisa Izaurralde (Addgene plasmid #146766; http://n2t.net/addgene:146766; RRID: Addgene_146766), and the following primer sequences: 5’-AACTTTCTGCCGATCCTCTTCAACCTGTATGGGCAGCCCGTGGCAGCCGGGGACACTCCAGCC-3’ and 5’-GGCTGGAGTGTCCCCGGCTGCCACGGGCTGCCCATACAGGTTGAAGAGGATCGGCAGAAAGTT-3’. The primer for sequencing is 5’-CGGCGGCTCTTGACCAGG-3’. Thereafter, the EGPF-tagged WT-RRP12 and RRP12Q699A plasmids were transfected into HCT 116 cells using Lipofectamine 2000 Transfection Reagent (Invitrogen) to overexpress the target proteins. The transfected cells were cultured without the exogenous treatment of serotonin and then lysed by using the lysis buffer containing 1% Triton X-100, 1X Tris Buffered Saline (TBS), and 1% protease inhibitor cocktail. 25 μL of ChromoTek GFP-Trap Magnetic Particles M-270 (Proteintech) were washed twice by using 1X PBS and then added to the protein supernatants and incubated with rotation for 1 hour at 4°C. Finally, the target protein-bound beads were collected and washed three times using the cell lysis buffer.

For the serotonylation analysis, the washed beads were resuspended in 100 μL of 100 mM phosphate buffer (pH 7). The probe labeling reaction was carried out on ice as described above. After the reaction, the supernatant was carefully removed, and the beads were washed three times using the phosphate buffer. The EGPF-tagged proteins were eluted by using 1X SDS Sample Loading Buffer (Sigma-Aldrich) at 95°C. The samples were cooled down to room temperature and subsequently subjected to SDS-PAGE and immunoblotting analyses. The EGFP-tagged proteins were imaged by using anti-GFP (CST, #2555) and the probe-labeled proteins were imaged by using IRDye 680RD Streptavidin (LI-COR).

### Data processing and statistical analysis

Fragpipe software (v21.1) was utilized for processing the MS and MS/MS raw data. The MS/MS spectra were aligned with the human UNIPROT database, which comprises 20,421 human reviewed entries. The applied search criteria included strict trypsin digestion and the incorporation of variable modifications such as serotonylated glutamine with or without the aryldiazonium-biotin probe modification (+707.3101 Da or +159.0684 Da), cysteine carboxyamidomethylation (+57.0214 Da), methionine oxidation (+15.9949 Da), and N-acylation (+42.0106 Da). The analysis allowed for up to three missed cleavages, and other search parameters were set to default values. For statistical analysis, Omicsbean software was employed. A gene filter was applied to calculate the fold-change values of protein expression. The t-test analysis was utilized to filter differentially expressed proteins, with a fold change threshold set at greater than 2-fold or less than 0.5-fold, and a significance level (P-value) below 0.05. Specifically, a two-sided t-test followed by Benjamini-Hochberg correction was applied to calculate the adjusted p values.

## Results and discussion

### Probe labeling and enrichment of serotonylated proteins from HCT 116 cells

The photoactive aryldiazonium-biotin probe (**Fig. 1**) was synthesized (**Supplementary Information**) and characterized, based on the protocols reported in our previous study.^14^ HCT 116 cells were cultured and utilized as a model of colorectal cancer to identify serotonylated proteins, where the experimental and control groups were treated with the same amount of serotonin to induce serotonylation on the target proteins. The cellular proteins were extracted and then treated with the UV-activated aryldiazonium-biotin probe (**Fig. 1**). Under the carefully controlled conditions of pH 6-7, the CRACR specifically occurs on the C4 position of 5-hydroxyindole moieties rather than C3 or C5 of tyrosine residues (**Fig. S1**). Therefore, the serotonylated proteins were labeled by the aryldiazonium-biotin probe and then enriched by using the streptavidin magnetic beads. The subsequent SDS analysis and silver staining indicated that serotonylated proteins were obviously and robustly enriched by using the aryldiazonium-biotin probe-based pull-down in the experimental groups (**Fig. S2**). Notably, due to the high serotonin concentration within FBS and cell culture media (∼350 nM), the exogenous feeding of serotonin to HCT 116 cells did not influence the enrichment of modified proteins using our probe (**Fig. S3**).

In a previous chemical proteomic study of serotonylation, 5-PT was used as an alkyne-tagged mimic of serotonin and treated to the cell lysates together with exogenous recombinant TGM2 enzyme,^18^ which was attributed to the poor cell-penetrating ability of 5-PT but might induce the artifact PTMs. In comparison with the 5-PT strategy, our aryldiazonium-biotin probe could sensitively capture the endogenous protein serotonylation from cultured cells and even tissues, thereby enabling the successful enrichment of a lot more modified proteins from cancer cells (**Fig. S2**).

### Global profiling of serotonylated proteome in HCT 116 cells

The collected raw MS data first underwent standardization for subsequent analyses (**Supplementary Data 1-3**). Distinct stratification among different groups was observed through Principal Components Analysis (PCA). The comparison analysis further indicated that there were 1,099 proteins significantly enriched in the aryldiazonium-biotin probe-treated group (**Fig. 2A**). Notably, these experiments were biologically repeated for at least three times and the data of enriched proteins were consistent (**Figs. 2B** and **S4**). Further signaling pathway analysis using the Kyoto Encyclopedia of Genes and Genomes (KEGG) database indicated that the enriched proteins possessing serotonylation were mainly located in the cell nucleus (**Fig. 2C**) and involved in more than ten pathophysiologically important pathways, which included amino acid biosynthesis, ribosome biogenesis, carbon metabolism, RNA degradation, *etc* (**Fig. 2E**). Notably, in line with dopaminylation, about 94% of the serotonylated proteins have the binding functions to proteins and RNA poly(A) tails (**Fig. 2D**). Biological Process (BP)-KEGG analysis further showed that the enriched proteins possessing serotonylation were mainly involved in ten basic but essential biological processes, including the regulation of gene expression, RNA processing, mRNA metabolism, peptide transportation, peptide biosynthesis, protein-complex assembly, *etc* (**Fig. 3A**). More importantly, utilizing the CRACR-based chemical proteomics approach, we successfully enriched and identified more proteins containing endogenous serotonylation than the previously reported 5-PT method by over twenty times (1,099:46), without inducing any artifacts. Even though different cell lines and probes were employed in these two studies, many serotonylated proteins identified by using 5-PT were included in the enriched proteins of this research (**Fig. S5**). Some of the serotonylated proteins [*e*.*g*., glyceraldehyde 3-phosphate dehydrogenase (GAPDH)] identified in this research have been verified by other studies.^23^ Finally, a number of overlapping proteins were observed between the serotonylated and dopaminylated proteome of HCT 116 cells (**Fig. S5**),^22^ confirming the enzymatic mechanism we proposed where TGM2 serves as an activation factor for the glutamine residues of the substrate proteins.^17^

**Figure 2.**
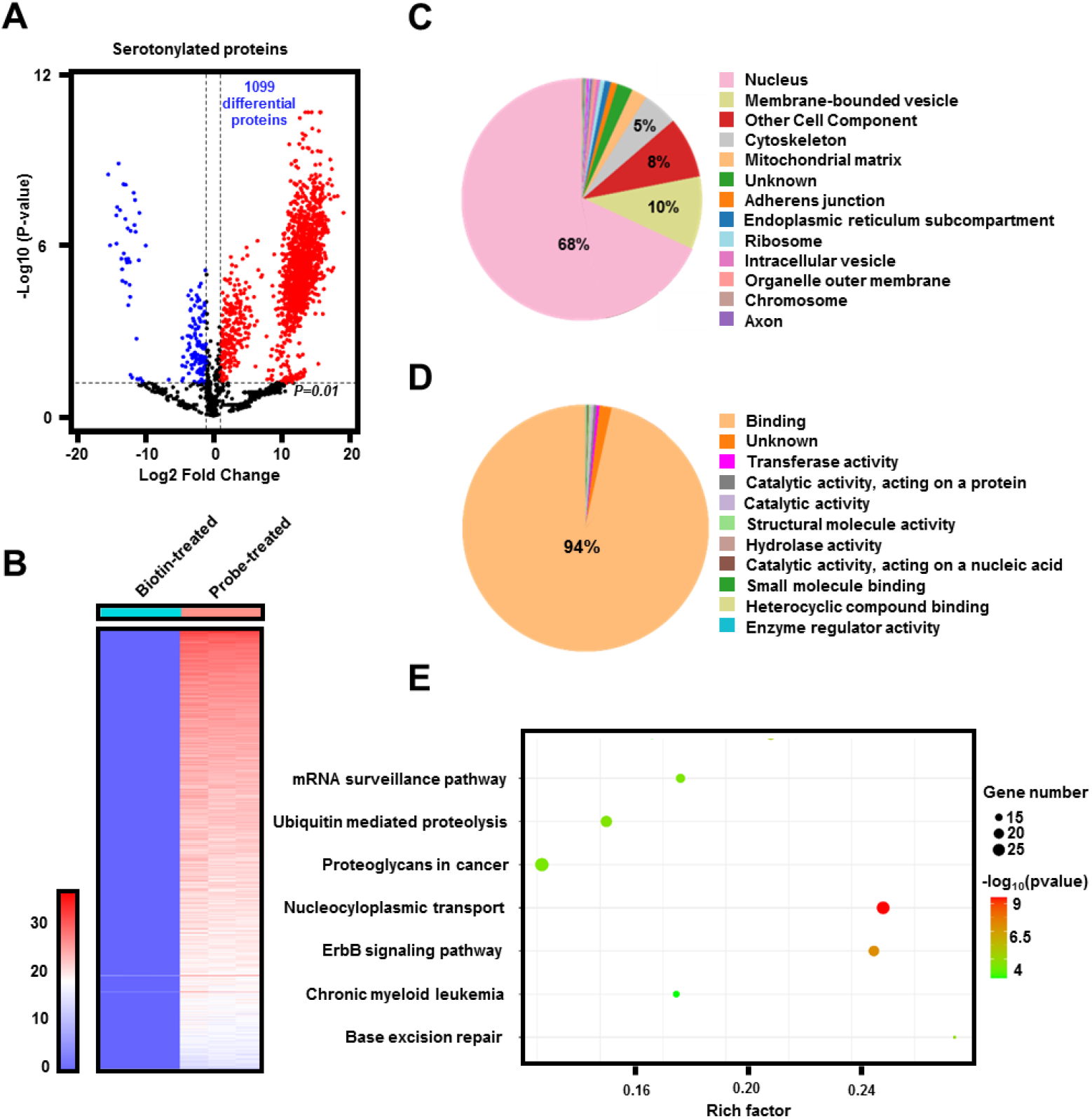
Global profiling of the serotonylation proteome in HCT 116 cells. (**A**) T-test comparison between the aryldiazonium-biotin probe-enriched and biotin-treated proteins. The analysis encompassed the data from three independent experiments. (**B**) Heatmap representation of label-free quantitation (LFQ) intensity data of the 1,099 serotonylated proteins enriched by using the aryldiazonium-biotin probe. The heatmap color (from red to blue) represents the fold change of protein level from increasing to decreasing. (**C**) Cell component of the identified proteins possessing serotonylation. (**D**) Molecular functions of the identified proteins possessing serotonylation. (**E**) KEGG (Kyoto Encyclopedia of Genes and Genomes) pathway analysis of the serotonylated proteins identified in this study.

**Figure 3.**
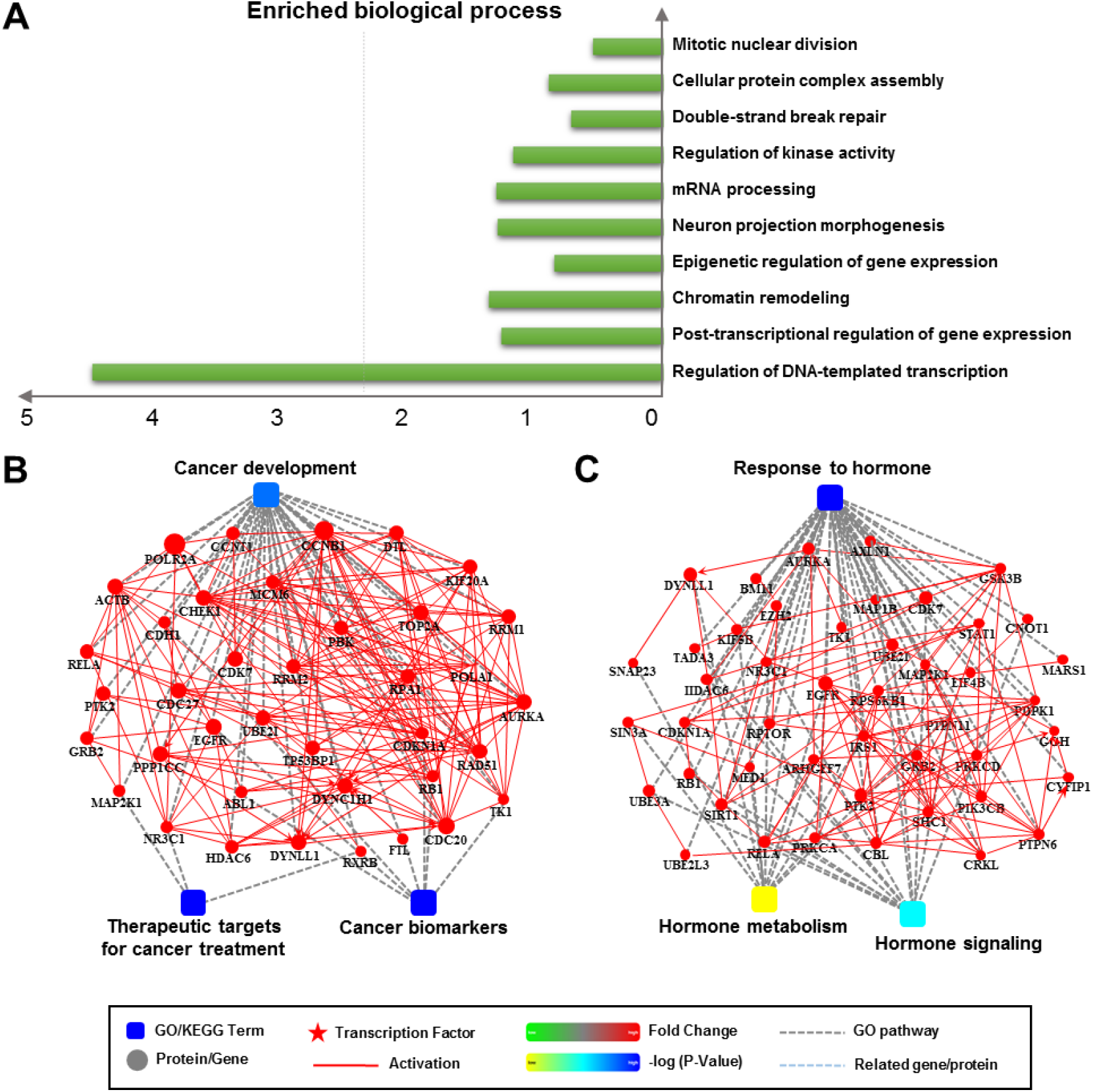
Function analysis of the serotonylation proteome in HCT 116 cells. (**A**) Top enriched biological processes that involve the serotonylated proteins identified in this study. (**B**) Interaction diagram illustrating the cluster of serotonylated proteins that are closely related to cancer biology. (**C**) Interaction diagram illustrating the cluster of serotonylated proteins that are closely related to hormone-based regulations. Color bar from green to red represents the fold change of protein levels from increasing to decreasing. The significance of the pathways represented by -log(p value) (Fisher’s exact test) was shown by color scales with dark blue as most significant.

### Interaction network analysis of enriched proteins containing serotonylation

To uncover the pathophysiological relevance of protein serotonylation, we conducted the interaction network analysis on the serotonylated proteins identified in this study and found that a number of them were associated with carcinogenesis and cancer development (**Fig. 3B**). Specifically, the interaction diagram illustrated that a number of serotonylated proteins were closely linked to the regulation of tumor necrosis factor (TNF), cancer metabolism, and cancer development/treatment, which included the mammalian target of rapamycin (mTOR),^24^ epidermal growth factor receptor (EGFR),^25^ 2’-5’-oligoadenylate synthetase (OAS3),^26^ rho-associated coiled-coil-containing protein kinase 1 (ROCK1),^27^ p21-activated kinase 4 (PAK4),^28^ histone deacetylase 6 (HDAC6),^29^ *etc*. The serotonylation on these proteins may exhibit significant regulatory functions for their activities. Notably, a variety of serotonylated proteins are common targets for cancer therapies (*e*.*g*., mTOR, EGFR, HDAC6),^24,25,29^ suggesting that to target the installation of serotonylation by TGM2 inhibitors may serve as a promising therapeutic strategy in clinic.

Notably, the protein interaction network analysis also illustrated that protein serotonylation was highly involved in the hormone-based regulations, including hormone biosynthesis and responses (**Fig. 3C**). These serotonylated proteins include ATPase Na^+^/K^+^ transporting subunit Alpha 1 (ATP1A1),^30^ thyroid hormone receptor associated protein 3 (THRAP3),^31^ EGFR,^32^ IQ motif-containing GTPase activating protein 1 (IQGAP1),^33^ 6-phosphofructokinase muscle type (PFKM),^34^ *etc*. These identified proteins possessing serotonylation exhibit key regulatory functions in hormone signaling pathways. The installation of an aromatic moiety, serotonin, onto the structurally important glutamine residues of these regulatory proteins may significantly influence their corresponding activities as transcription factors, transporters, receptors, or enzymes (**Fig. 3C**). Overall, our results in this chemical proteomic study will open a new door for understanding the pathological relevance of serotonin *via* its covalent modifications on regulatory proteins.

### Identification of serotonylation sites

Unlike protein dopaminylation that was found on both glutamine and cystine,^22^ serotonylation can only occur on glutamine residues *via* transamination.^35^ The C4 position of 5-hydroxyindole is electron-rich and can be selectively modified by our aryldiazonium-biotin probe *via* pH-controlled CRACR.^19^ Therefore, the aryldiazonium-biotin probe-tagged proteins containing serotonylation could be enriched and identified by mass spectrometry. As the azo structure formed between the probe and 5-hydroxyindole can be cleaved under the ionization conditions of mass spectrometer, there are two possible mass shifts in the modified peptide fragments (**Fig. 4A**). The serotonylation sites of many key regulatory proteins, such as myosin-9 (MYH9),^36^ plectin (PLEC),^37^ ribosomal RNA processing 12 homolog (RRP12),^38^ ubiquitin protein ligase E3 component N-recognin 4 (UBR4),^39^ *etc*, were identified based on the high-quality mass spectra (**Fig. 4B** and **Table S1**). Notably, the two possible mass shifts (+707.3101 Da and +159.0684 Da) were both observed on the same peptide fragment of RRP12 and the modification site was Q699 (**Fig. 4B**), which strongly proved the accuracy and sensitivity of our chemical proteomics approach for profiling protein serotonylation. The site-directed mutagenesis and immunoblotting analysis using our biotin-tagged probe further confirmed the Q699 residue of RRP12 was indeed modified by serotonin (**Figs. 4C** and **S6**). In general, the serotonylated sites identified via our chemical proteomic approach may significantly influence the activities and functions of these pathophysiologically important proteins through the allosteric effect, steric effect, electronic effect, π-π stacking, *etc*.^14^ Notably, serotonylation may have crosstalk with other protein PTMs (such as ubiquitination and acetylation) by directly influencing the enzymatic activities of the corresponding writers (*e*.*g*., UBR4)^40^ or erasers (*e*.*g*., HDAC6)^29^.

**Figure 4.**
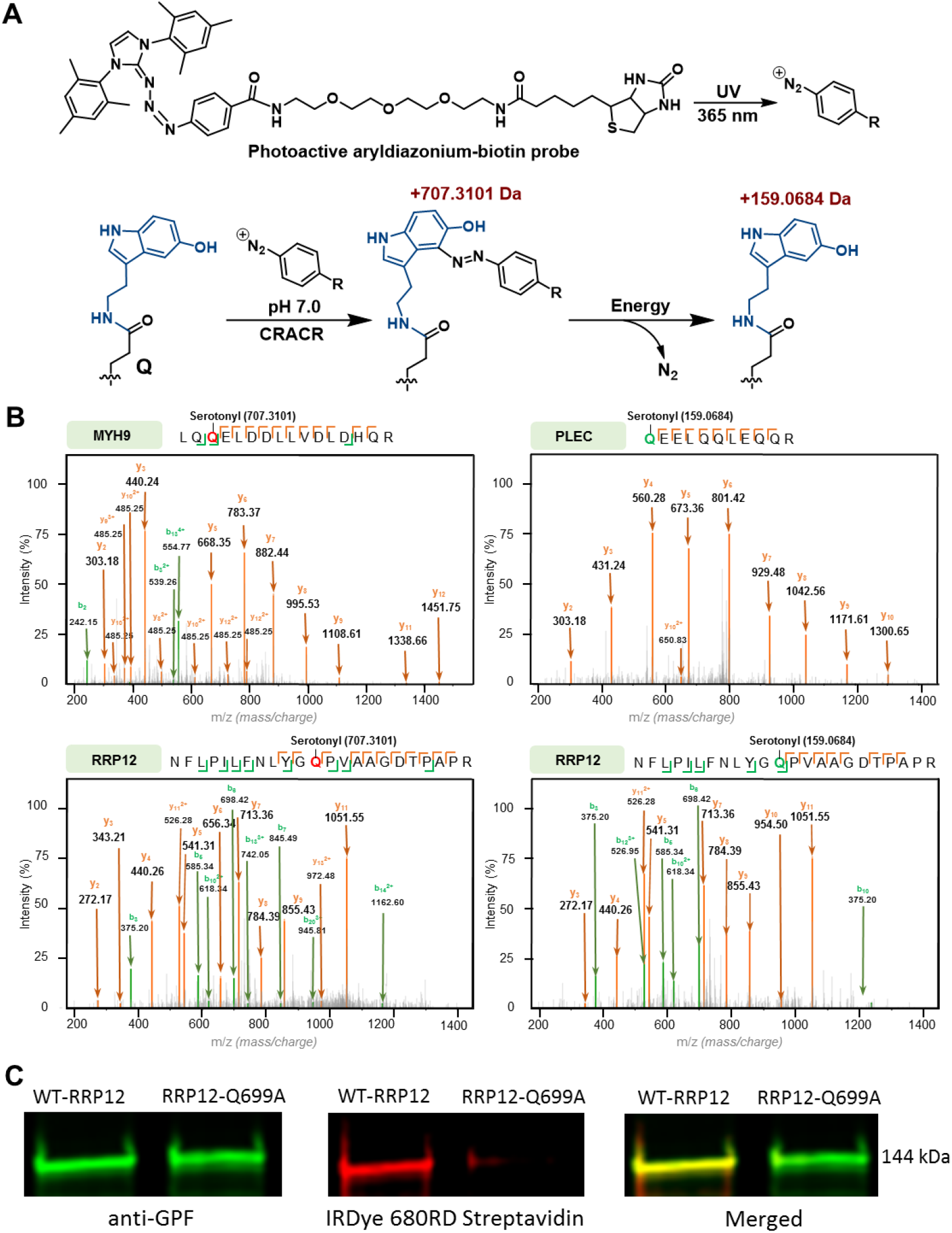
Modification site analysis of the enriched proteins possessing serotonylation in HCT 116 cells. (**A**) Fragment structures of the serotonylated proteins and the corresponding mass shifts. (**B**) Examples of the identified modification sites and the corresponding mass shifts. (**C**) Biochemical validation of RPP12 serotonylation on its Q699 residue by using site-directed mutagenesis, immunoprecipitation, and immunoblotting.

## Conclusion

Serotonin is a monoamine metabolite biosynthesized from the essential amino acid, tryptophan,^41^ which plays key regulatory roles in cell signaling as a well-known neurotransmitter.^42^ The discovery of serotonylation on protein glutamine residues provided a novel mode of action (MOA) to serotonin-mediated signaling pathways. One well-established example is that histone H3 serotonylation (H3Q5ser) has been shown to regulate gene transcription by enhancing TFIID binding to H3K4me3 and stabilizing the epigenetic mark H3K4me3.^10-14^ In our previous studies, we have uncovered that the dynamics of histone H3 serotonylation is regulated by the single enzyme, TGM2.^17^ Because of the enzyme promiscuity of transglutaminases, there are diverse non-histone proteins individually reported undergoing serotonylation on glutamine residues.^5^ However, due to the limitations of research tools and methodologies, the global profiling of protein serotonylation still remains challenging and the full landscape of serotonylated proteome is poorly understood.

Recently, we have applied an aryldiazonium-biotin probe to label and enrich histone serotonylation in cultured cell and collected tissue samples in a biorthogonal manner. This efficient chemical biology approach relays on a pH-controlled azo-coupling reaction, *i*.*e*., CRACR.^19^ Utilizing this powerful tool, we discovered that histone serotonylation was accumulated in tumor cells that overexpress TGM2 (*e*.*g*., colon and breast cancer cells) and played significant roles in regulating the cellular chromatin structure.^14^ In this research, we employed this aryldiazonium-biotin probe to label and enrich the serotonylated proteome in colorectal cancer (HCT 116) cells for a MS-based global proteomic profiling. Thus, 1,099 proteins possessing serotonylation were identified, which are mainly located in the cell nucleus and involved in a variety of basic but essential signaling pathways. Further bioinformatic analyses revealed that these serotonylated proteins were primarily related to cancer development/treatment and hormone-based regulations. The highly overlapping proteins from serotonylated and dopaminylated proteome of HCT 116 cells proved solid evidence for the enzymatic mechanism we proposed in our previous research, where TGM2 acts as an activation factor by forming a thioester structure with the substrate. Finally, utilizing the chemical proteomics approach developed in this study, we have successfully identified a number of serotonylation sites from various pathophysiologically important proteins. Altogether, we utilized a novel aryldiazonium-biotin probe to conduct the global profiling of protein serotonylation in colon cancer cells and identified over 1,000 serotonylated proteins that are closely associated with cancer development and treatment. The dataset generated by using our new methodology in this work highlighted the importance of serotonylation in cell signaling and gave new insights for novel disease therapeutics in the future.

## Data Availability

The mass spectrometry proteomics data have been deposited to the ProteomeXchange Consortium (https://proteomecentral.proteomexchange.org) via the iProX partner repository with the data set identifier PXD054426.

## Supporting Information

The Supporting Information is available free of charge at https://pubs.acs.org/doi/10.1021/acs.jproteome.XXXX.

General methods (equipment, reagents, chemicals); synthesis of the photoactive aryldiazonium-biotin probe; statistics and reproducibility; serotoninylation site analysis (Table S1); reaction selectivity of pH-controlled CRACR on protein residues (Figure S1); SDS-PAGE and silver staining analysis of enriched proteins possessing serotoninylation (Figure S2); silver staining analysis of the probe-based enrichment of serotonylated proteins from HCT 116 cells with and without serotonin treatment (Figure S3); distribution map of the data in each sample (Figure S4); Venn diagram analysis of the overlapping proteins between the serotonylated, dopaminylated, and 5-PT-modified proteome (Figure S5); uncropped immunoblotting images (Figure S6) (PDF)

Raw data from mass spectrometry-based quantitative proteomic analysis (Data S1) (XLSX)

Identified proteins with serotonylation (Data S2) (XLSX)

KEGG analysis (Data S3) (XLSX)

Enriched biological processes (Data S4) (XLSX)

Peptides with target PTMs (Data S5) (XLSX)

## Notes

The authors declare that they have no known competing financial interests or personal relationships that could have appeared to influence the work reported in this paper.

## Acknowledgements

We acknowledge Drs. Liwen Zhang and Sophie Harvey and the Campus Chemical Instrument Center (CCIC) Mass Spectrometry and Proteomics Facility at OSU for their assistance in mass spectroscopy.

## Reference

1. Sarkar N. K.; Clarke D. D.; Waelsch H. An enzymically catalyzed incorporation of amines into proteins. Biochim. Biophys. Acta 1957, 25, 451.

2. Clarke D. D.; Mycek M. J.; Neidle A.; Waelsch H. The incorporation of amines into protein. Arch. Biochem. Biophys. 1959, 79, 338.

3. Gundemir, S.; Colak, G.; Tucholski, J.; Johnson, G. V. Transglutaminase 2: A molecular Swiss army knife. Biochim. Biophys. Acta. 2012, 1823, 406.

4. Hummerich, R.; Thumfart, J. O.; Findeisen, P.; Bartsch, D.; Schloss, P. Transglutaminase-mediated transamidation of serotonin, dopamine and noradrenaline to fibronectin: Evidence for a general mechanism of monoaminylation. FEBS Lett. 2012, 586, 3421.

5. Bader, M. Serotonylation: Serotonin signaling and epigenetics. Front. Mol. Neurosci. 2019, 12, 288.

6. Walther, D. J.; Peter, J. U.; Winter, S.; Höltje, M.; Paulmann, N.; Grohmann, M.; Vowinckel, J.; Alamo-Bethencourt, V.; Wilhelm, C. S.; Ahnert-Hilger, G.; Bader, M. Serotonylation of small GTPases is a signal transduction pathway that triggers platelet alpha-granule release. Cell 2003, 115, 851.

7. Dale, G. L.; Friese, P.; Batar, P.; Hamilton, S. F.; Reed, G. L.; Jackson, K. W.; Clemetson, K. J.; Alberio, L. Stimulated platelets use serotonin to enhance their retention of procoagulant proteins on the cell surface. Nature 2002, 415, 175.

8. Hummerich, R.; Costina, V.; Findeisen, P.; Schloss, P. Monoaminylation of fibrinogen and glia-derived proteins: Indication for similar mechanisms in posttranslational protein modification in blood and brain. ACS Chem. Neurosci. 2015, 6, 1130.

9. Cooper, A.; Woulfe, D.; Kilic, F. Post-translational modifications of serotonin transporter. Pharmacol. Res. 2019, 140, 7.

10. Farrelly, L. A.; Thompson, R. E.; Zhao, S.; Lepack, A. E.; Lyu, Y.; Bhanu, N. V.; Zhang, B.; Loh, Y. E.; Ramakrishnan, A.; Vadodaria, K. C.; Heard, K. J.; Erikson, G.; Nakadai, T.; Bastle, R. M.; Lukasak, B. J.; Zebroski, H. 3rd; Alenina, N.; Bader, M.; Berton, O.; Roeder, R. G.; Molina, H.; Gage, F. H.; Shen, L.; Garcia, B. A.; Li, H.; Muir, T. W.; Maze, I. Histone serotonylation is a permissive modification that enhances TFIID binding to H3K4me3. Nature 2019, 567, 535.

11. Zlotorynski, E. Histone serotonylation boosts neuronal transcription. Nat. Rev. Mol. Cell Biol. 2019, 20, 323.

12. Zhao, S.; Chuh, K. N.; Zhang, B.; Dul, B. E.; Thompson, R. E.; Farrelly, L. A.; Liu, X.; Xu, N.; Xue, Y.; Roeder, R. G.; Maze, I.; Muir, T. W.; Li, H. Histone H3Q5 serotonylation stabilizes H3K4 methylation and potentiates its readout. Proc. Natl. Acad. Sci. U. S. A. 2021, 118, e2016742118.

13. Zhao, J.; Chen, W.; Pan, Y.; Zhang, Y.; Sun, H.; Wang, H.; Yang, F.; Liu, Y.; Shen, N.; Zhang, X.; Mo, X.; Zang, J. Structural insights into the recognition of histone H3Q5 serotonylation by WDR5. Sci. Adv. 2021, 7, eabf4291.

14. Zhang, N.; Wu, J.; Hossain, F.; Peng, H.; Li, H.; Gibson, C.; Chen, M.; Zhang, H.; Gao, S.; Zheng, X.; Wang, Y.; Zhu, J.; Wang, J. J.; Maze, I.; Zheng, Q. Bioorthogonal labeling and enrichment of histone monoaminylation reveal its accumulation and regulatory function in cancer cell chromatin. J. Am. Chem. Soc. 2024, 146, 16714–16720.

15. Zhang, N.; Wu, J.; Zheng, Q. Chemical proteomics approaches for protein post-translational modification studies. Biochim. Biophys. Acta Proteins Proteom. 2024, 1872, 141017.

16. Lin, J. C.; Chou, C. C.; Gao, S.; Wu, S. C.; Khoo, K. H.; Lin, C. H. An in vivo tagging method reveals that Ras undergoes sustained activation upon transglutaminase-mediated protein serotonylation. Chembiochem 2013, 14, 813.

17. Zheng, Q.; Bastle, R. M.; Zhao, S.; Kong, L.; Vostal, L.; Ramakrishnan, A.; Shen, L.; Fulton, S. L.; Wang, H.; Zhang, B.; Upad, A.; Dierdorff, L.; Thompson, R. E.; Molina, H.; Stransky, S.; Sidoli, S.; Muir, T. W.; Li, H.; David, Y.; Maze, I. Histone monoaminylation dynamics are regulated by a single enzyme and promote neural rhythmicity. bioRxiv 2022-12-06, doi: 10.1101/2022.12.06.519310.

18. Lin, J. C.; Chou, C. C.; Tu, Z.; Yeh, L. F.; Wu, S. C.; Khoo, K. H.; Lin C. H. Characterization of protein serotonylation via bioorthogonal labeling and enrichment. J. Proteome. Res. 2014, 13, 3523.

19. Addy, P. S.; Erickson, S. B.; Italia, J. S.; Chatterjee, A. A chemoselective rapid azo-coupling reaction (CRACR) for unclickable bioconjugation. J. Am. Chem. Soc. 2017, 139, 11670.

20. Guzmán, L. E.; Wijetunge, A. N.; Riske, B. F.; Massani, B. B.; Riehle, M. A.; Jewett, J. C. Chemical probes to interrogate the extreme environment of mosquito larval guts. J. Am. Chem. Soc. 2024, 146, 8480.

21. Bertolini, M.; Mendive-Tapia, L.; Ghashghaei, O.; Reese, A.; Lochenie, C.; Schoepf, A. M.; Sintes, M.; Tokarczyk, K.; Nare, Z.; Scott, A. D.; Knight, S. R.; Aithal, A. R.; Sachdeva, A.; Lavilla, R.; Vendrell, M. Nonperturbative fluorogenic labeling of immunophilins enables the wash-free detection of immunosuppressants. ACS Cent. Sci. 2024, 10, 969–977.

22. Zhang, N.; Gao, S.; Peng, H.; Wu, J.; Li, H.; Gibson, C.; Wu, S.; Zhu, J.; Zheng, Q. Chemical proteomic profiling of protein dopaminylation in colorectal cancer cells. J. Proteome Res. 2024, 23, 2651–2660.

23. Wang, X.; Fu, S. Q.; Yuan, X.; Yu, F.; Ji, Q.; Tang, H. W.; Li, R. K.; Huang, S.; Huang, P. Q.; Qin, W. T.; Zuo, H.; Du, C.; Yao, L. L.; Li, H.; Li, J.; Li, D. X.; Yang, Y.; Xiao, S. Y.; Tulamaiti, A.; Wang, X. F.; Dai, C. H.; Zhang, X.; Jiang, S. H.; Hu, L. P.; Zhang, X. L.; Zhang, Z. G. Mol. Cell. 2024, 84, 760–775.

24. Tian, T.; Li, X.; Zhang, J. mTOR signaling in cancer and mTOR inhibitors in solid tumor targeting therapy. Int. J. Mol. Sci. 2019, 20, 755.

25. Uribe, M. L.; Marrocco, I.; Yarden, Y. EGFR in cancer: Signaling mechanisms, drugs, and acquired resistance. Cancers 2021, 13, 2748.

26. Li, X. Y.; Hou, L.; Zhang, L. Y.; Zhang, L.; Wang, D.; Wang, Z.; Wen, M. Z.; Yang, X. T. OAS3 is a co-immune biomarker associated with tumour microenvironment, disease staging, prognosis, and treatment response in multiple cancer types. Front. Cell Dev. Biol. 2022, 10, 815480.

27. Kim, S.; Kim, S. A.; Han, J.; Kim, I. S. Rho-kinase as a target for cancer therapy and its immunotherapeutic potential. Int. J. Mol. Sci. 2021, 22, 12916.

28. Yuan, Y.; Zhang, H.; Li, D.; Li, Y.; Lin, F.; Wang, Y.; Song, H.; Liu, X.; Li, F.; Zhang, J. PAK4 in cancer development: Emerging player and therapeutic opportunities. Cancer Lett. 2022, 545, 215813.

29. Aldana-Masangkay, G. I.; Sakamoto, K. M. The role of HDAC6 in cancer. J. Biomed. Biotechnol. 2011, 2011, 875824.

30. Gomez-Sanchez, C. E.; Kuppusamy, M.; Gomez-Sanchez, E. P. Somatic mutations of the ATP1A1 gene and aldosterone-producing adenomas. Mol. Cell Endocrinol. 2015, 408, 213.

31. Wang, Y. P.; Ma, C.; Yang, X. K.; Zhang, N.; Sun, Z. G. Pan-cancer and single-cell analysis reveal THRAP3 as a prognostic and immunological biomarker for multiple cancer types. Front Genet. 2024, 15, 1277541.

32. Herbst, R. S. Review of epidermal growth factor receptor biology. Int. J. Radiat. Oncol. Biol. Phys. 2004, 59, 21.

33. Abel, A. M.; Schuldt, K. M.; Rajasekaran, K.; Hwang, D.; Riese, M. J.; Rao, S.; Thakar, M. S.; Malarkannan, S. IQGAP1: insights into the function of a molecular puppeteer. Mol. Immunol. 2015, 65, 336.

34. Cen, L. P.; Ng, T. K.; Chu, W. K.; Pang, C. P. Growth hormone-releasing hormone receptor signaling in experimental ocular inflammation and neuroprotection. Neural Regen. Res. 2022, 17, 2643.

35. Muma, N. A.; Mi, Z. Serotonylation and transamidation of other monoamines. ACS Chem. Neurosci. 2015, 6, 961.

36. Pecci, A.; Ma, X.; Savoia, A.; Adelstein, R. S. MYH9: Structure, functions and role of non-muscle myosin IIA in human disease. Gene 2018, 664, 152.

37. Perez, S. M.; Brinton, L. T.; Kelly, K. A. Plectin in cancer: From biomarker to therapeutic target. Cells 2021, 10, 2246.

38. An, G.; Liu, Y.; Hou, Y.; Lei, Y.; Bai, J.; He, L.; Liu, Y. RRP12 suppresses cell migration and invasion in colorectal cancer cell via regulation of epithelial-mesenchymal transition. J. Gastrointest. Oncol. 2023, 14, 2111.

39. Jeong, D. E.; Lee, H. S.; Ku, B.; Kim, C. H.; Kim, S. J.; Shin, H. C. Insights into the recognition mechanism in the UBR box of UBR4 for its specific substrates. Commun Biol. 2023, 6, 1214.

40. Barnsby-Greer, L.; Mabbitt, P. D.; Dery, M. A.; Squair, D. R.; Wood, N. T.; Lamoliatte, F.; Lange, S. M.; Virdee, S. UBE2A and UBE2B are recruited by an atypical E3 ligase module in UBR4. Nat. Struct. Mol. Biol. 2024, 31, 351.

41. Perez-Castro, L.; Garcia, R.; Venkateswaran, N.; Barnes, S.; Conacci-Sorrell, M. Tryptophan and its metabolites in normal physiology and cancer etiology. FEBS J. 2023, 290, 7.

42. Yabut, J. M.; Crane, J. D.; Green, A. E.; Keating, D. J.; Khan, W. I.; Steinberg, G. R. Emerging roles for serotonin in regulating metabolism: New implications for an ancient molecule. Endocr. Rev. 2019, 40, 1092.

