## Supplemental Information for "pH-Controlled chemoselective rapid azo-coupling reaction (CRACR) enables global profiling of serotonylation proteome in cancer cells"

### Table of Contents

**Materials and Methods**------------------------------------------------------------------------**3**

**Table S1.** Modification site analysis-------------------------------------------------------------**5**

**Figure S1.** Reaction selectivity of pH-controlled CRACR on protein residues---------**6**

**Figure S2.** SDS-PAGE analysis of the enriched proteins possessing serotonylation---**7**

**Figure S3.** Comparison of the serotonylation proteome from cells cultured with and without exogenous serotonin-----------------------------------------------------**8**

**Figure S4.** Distribution map of the data in each sample ----------------------------------**9**

**Figure S5.** Venn diagram analysis of the overlapping proteins between the serotonylated and dopaminylated proteome--------------------------------**10**

**Figure S6.** Uncropped immunoblotting images--------------------------------------------**11**

**References and Notes**-------------------------------------------------------------------------**12**

**Data S1.** Raw data from mass spectrometry-based quantitative proteomic analysis.

**Data S2.** Identified proteins with serotonylation.

**Data S3.** KEGG analysis.

**Data S4.** Enriched biological processes.

**Data S5.** Peptides with target PTMs.

### MATERIALS AND METHODS

#### General methods (equipment, reagents, chemicals)

UV spectrometry was performed on a NanoDrop 2000c (Thermo Scientific). Biochemicals and media were purchased from Fisher Scientific or Sigma-Aldrich Corporation unless otherwise stated. Centrifugal filtration units were purchased from Millipore, and MINI dialysis units purchased from Pierce. Gels were imaged on an Odyssey CLx Imaging System (Li-Cor). The primary antibodies were purchased from Cell Signaling Technology (CST) and the secondary antibodies were purchased from Li-Cor. The primers were from Integrated DNA Technologies (IDT). All the experiments in this research were performed at least 3X. For the synthesis of probe molecules used in this study, all commercial chemicals were purchased from Sigma Aldrich, TCI chemicals, AK Scientific, Fischer Scientific, Broadpharm and used without further purification. The reagents and solvents were handled following the safety processes required as instructed by the manufacturer. Organic and aqueous waste was disposed of following standard safety protocols. Solvents for workup were purchased from Fisher Chemical, and anhydrous solvents were purchased from Sigma Millipore in a sealed bottle and degassed by passing $N_2$ before each use. Reaction progress was monitored by using normal phase TLC silica gel 60 $F_{254}$ plates by Sigma Aldrich (aluminum backed 20 X 20 cm). Developed plates were analyzed by visualizing under a UV-light and/or staining with Phosphomolybdic Acid (PMA) Stain (100 mL absolute ethanol and 10 g PMA). Isolation and purification of the crude reaction materials were performed using silica gel ($SiO_2$) by Acros Organic (0.030-0.200 mm, 60 $A^0$). Organic solvents were removed under vacuum using a Heidolph Rotavapor equipped with a dry ice condenser.

#### Synthesis of the photoactive aryldiazonium-biotin probe

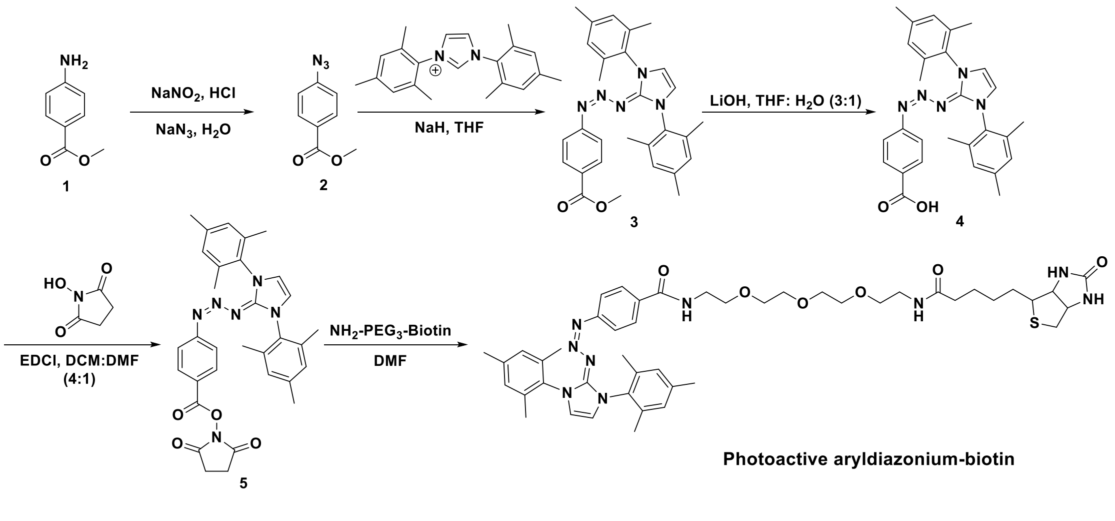


The photoactive aryldiazonium-biotin probe was synthesized based on the protocols reported in our previous study.[1] Briefly, the amino group of starting material, methyl-4-amino benzoate (**1**), was converted to the azido group in the presence of acidic solutions of sodium nitrite ($NaNO_2$) and sodium azide ($NaN_3$). Thereafter, a trizabutadiene group was installed to methyl 4-azidobenzoate (**2**) utilizing 1,3-dimesitylimidazolium chloride and sodium hydride (NaH). The ester group of **3** was then deprotected with aqueous lithium hydroxide (LiOH) to generate compound **4**. The carboxylic acid group of **4** was subsequently activated with *N*-hydroxysuccinamide and 1-(3-Dimethylaminopropyl)-3-ethylcarbodiimide hydrochloride hydrochloride (EDCl) to give **5**. At last, compound **5** was conjugated with biotin-PEG$_3$-amine to get the photoactive probe. The aryldiazonium-biotin probe was purified through recrystallization for the NMR analysis[1] and described applications in this study.

**Statistics and reproducibility**

All the mass spectrometry analyses were repeated independently at least three times with similar results. Significance was determined at $p<0.05$. All data are represented as mean ± SEM. Statistical analyses were performed in GraphPad Prism 9.

### SUPPLEMENTARY TABLES

| Peptide Sequence | Modified Sites | Gene Name |
|---|---|---|
| QAMLENASDIKLEK | 1Q(159.0684) | ABCF1 |
| TNIQNLNQLREDELGSDISALTLR | 4Q(707.3101) | AKAP9 |
| AGAAPVAPEKQATELALLQR | 11Q(159.0684) | ARHGEF2 |
| QAADMILLDDNFASIVTGVEEGR | 1Q(159.0684) | ATP1A1 |
| IHISQEDNVANKQTLASYR | 13Q(159.0684) | BAZ1A |
| QVTESSHLEQQLEEANAVR | 1Q(159.0684) | CDC42BPA |
| TDAKQKWLLLTGISAQQNR | 5Q(159.0684) | CLTC |
| IKQLIELDYLNPGSIR | 3Q(159.0684) | DDX20 |
| GVTFLFPIQAKTFHHVYSGK | 9Q(159.0684) | DDX21 |
| QYPISLVLAPTR | 1Q(159.0684) | DDX3X |
| NLGIESQDVMQQATNAILR | 10M(15.9949),7Q(159.0684) | DDX46 |
| QYLPIFAVQQELLTIIR | 1Q(159.0684) | DHX38 |
| DVTNNQEKHFYTFTNEHHR | 6Q(159.0684) | DIS3 |
| LQMEAPHIIVGTPGR | 2Q(159.0684) | EIF4A1 |
| QEEFKHIAVFPCK | 12C(57.0215),1Q(159.0684) | EIF5B |
| FDASFFGVHPKQAHTMDPQLR | 12Q(159.0684) | FASN |
| LFDHPESPTPNPTEPLFLAQAEVYK | 20Q(159.0684) | FASN |
| QKLYTLQDKAQVADVVVSR | 7Q(159.0684) | FASN |
| QNAPMTLEEFRK | 1Q(159.0684) | GBF1 |
| QADFEAHNILR | 1Q(159.0684) | GMPS |
| ALSAVSTQQKKADR | 9Q(159.0684) | GOLGA2 |
| SSVAVLTQESFAEHR | N-term(42.0106),8Q(707.3101) | HDLBP |
| QHYVLAGASGSPGEEVAIRPSTAPR | 1Q(159.0684) | HIP1R |
| QLQDIATLADQR | 1Q(159.0684) | IPO7 |
| QIPAITCIQSQWR | 1Q(159.0684),7C(57.0215) | IQGAP1 |
| QFSVPPAPTRPSCPAVAEIPLR | 13C(57.0215),1Q(159.0684) | KIF2C |
| LSTQEGLVQELQKKQVELQEER | 9Q(159.0684) | KIFC1 |
| ELNELKDQIQDVEGKYMQGLK | 17M(15.9949),8Q(159.0684) | LRRFIP1 |
| SLLQAWGLILR | 4Q(159.0684) | MDN1 |
| MLQTLKELEAHSAEAR | 1M(15.9949),3Q(159.0684) | MYBBP1A |
| QELELVVSELEAR | 1Q(159.0684) | MYH14 |
| KKMQQNIQELEEQLEEEESAR | 3M(15.9949),4Q(707.3101),5Q(159.0684) | MYH9 |
| LQQELDDLLVDLDHQR | 3Q(707.3101) | MYH9 |
| QELEEIC[57.0215]HDLEAR | 1Q(159.0684),7C(57.0215) | MYH9 |
| QKHSQAVEELAEQLEQTKR | 1Q(159.0684) | MYH9 |
| QDDPFELFIAATNIR | 1Q(707.3101) | NAT10 |
| LVMAESEKSQKLEER | 10Q(159.0684) | NUMA1 |
| QQNQELQEQLR | 1Q(159.0684) | NUMA1 |
| PNPQVYSR | 4Q(159.0684) | OAS3 |
| QEELQQLEQQR | 1Q(159.0684) | PLEC |
| QQIAEIEKQTKEQSQLTATQTR | 9Q(159.0684) | PRPF8 |
| WSKQTDVGITHFR | 4Q(159.0684) | PRPF8 |
| AQAALQAVNSVQSGNLALAASAAAVDAGMAMAGQSPVLR | 12Q(159.0684) | PTBP1 |
| QLPPPPPPIPPPRPLIQR | 1Q(159.0684) | RBBP6 |
| QSAQFWNATFAK | 1Q(159.0684) | RIF1 |
| RDSFLAQTKNK | 7Q(159.0684) | RIF1 |
| QVVNIPSFIVR | 1Q(159.0684) | RPS9 |
| NFLPILFNLYGQPVAAGDTPAPR | 12Q(707.3101) | RRP12 |
| NFLPILFNLYGQPVAAGDTPAPR | 12Q(159.0684) | RRP12 |
| SSLGQSASETEEDTVSVSKK | 5Q(159.0684) | SF3B2 |
| EYLEGSNLITKLQAKHDLLQR | 13Q(159.0684) | SRGAP1 |
| QFVQSAKEVANSTANLVK | 4Q(159.0684) | TLN1 |
| TYNASITLQQQLKELTAPDENIPAK | 11Q(159.0684) | TOP1 |
| LQEKLSPPYSSPQEFAQDVGR | 2Q(159.0684) | TRIM28 |
| QMLFYVTAFDRDR | 1Q(159.0684) | TRIP12 |
| QGLVNVALDILSR | 1Q(159.0684) | TRRAP |
| PGQSFQEQVEHYR | 3Q(159.0684) | U2SURP |
| QQGYSTVSHFNIVHYDHLAAVR | 17C(57.0215),1Q(159.0684) | UBR4 |
| TAEQSCDQKLTNTNR | 6C(57.0215),8Q(159.0684) | UHRF1 |
| QEVQAWDGEVR | 1Q(159.0684) | USP5 |
| AKEGQKADFPAGIPECGTDALR | 16C(57.0215),5Q(159.0684) | VARS |
| TVLDLIVEDLQSTSEDKEKQYTSQTTR | 11Q(159.0684) | VIRMA |
| QILHGDPLPLTR | 1Q(159.0684) | WDHD1 |

The serotoninylated glutamine (Q) residues are marked in red.

**Table S1.** Verified serotoninylation sites and the corresponding peptide sequences.

### SUPPLEMENTARY FIGURES AND LEGENDS

**Figure S1.** Reaction selectivity of pH-controlled CRACR on protein residues.

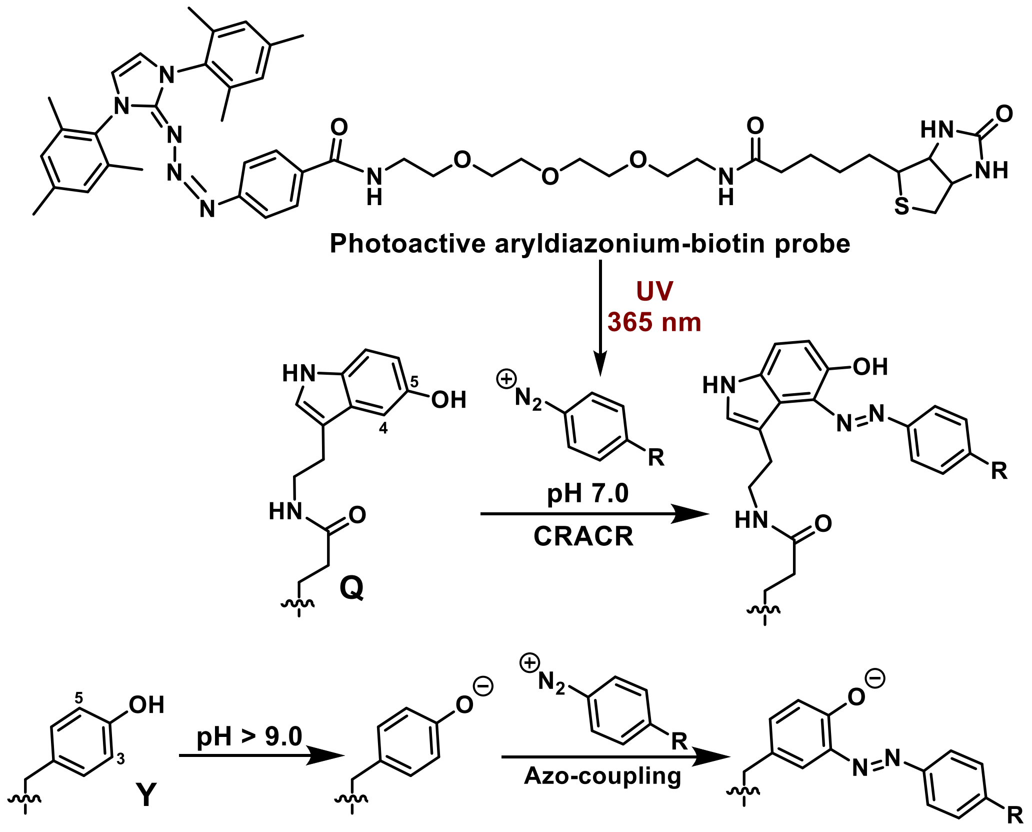

**Figure S2.** Aryldiazonium-biotin probe-based enrichment of serotonylated proteins from HCT 116 cells, indicating obvious enriched protein bands in the probe-treated groups.

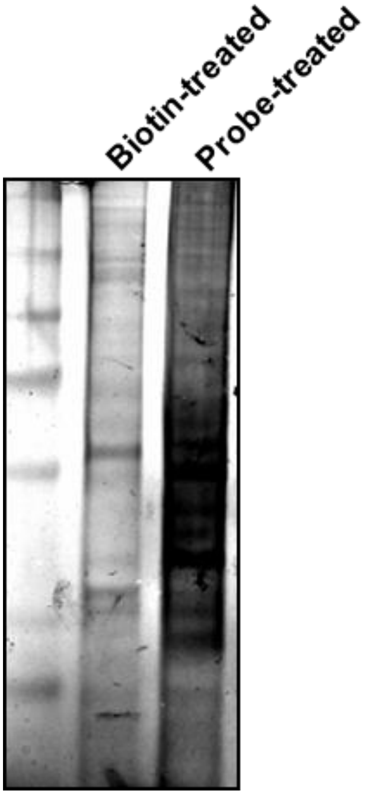

**Figure S3.** Silver staining analysis of the probe-based enrichment of serotonylated proteins from HCT 116 cells with and without serotonin treatment, illustrating that the exogenous feeding of serotonin to cultured cells did not significantly influence the serotonylation proteome.

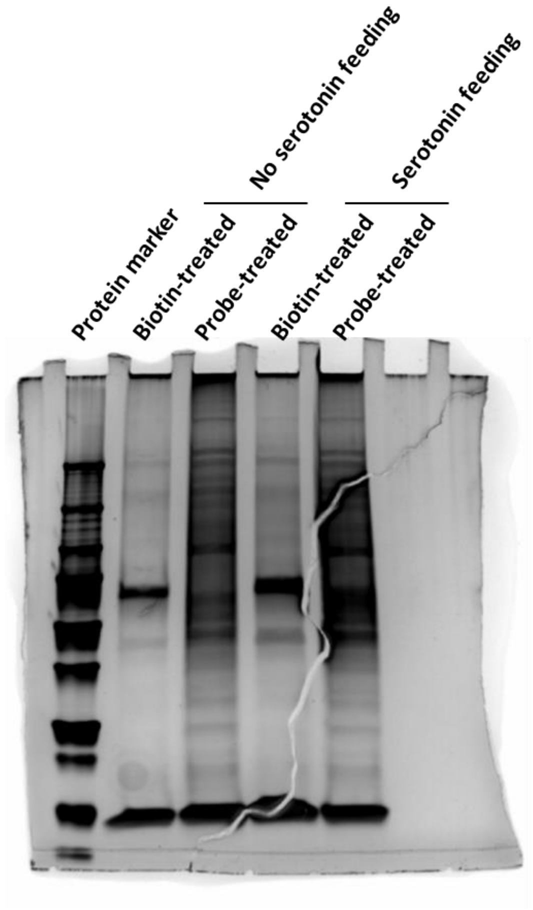

**Figure S4.** Distribution map of the data in each sample, *i.e.*, experimental groups (E1, E2 and E3) and control groups (C1, C2 and C3) before and after standardization.

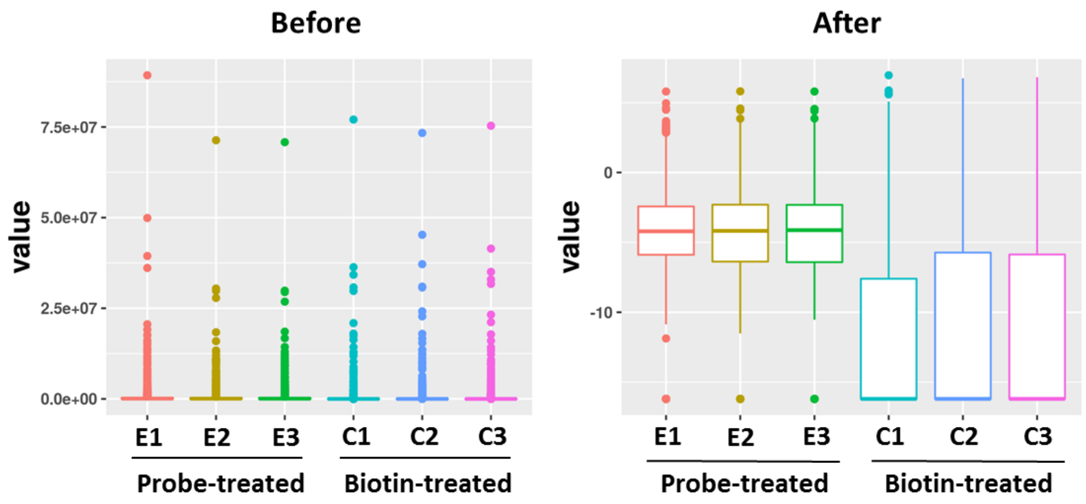

**Figure S5.** Venn diagram illustrating the overlapping proteins between the serotonylated (**A**) and dopaminylated (**B**) proteome[2] identified from HCT 116 cells and 5-PT-modified proteome identified from SW480 cells (**C**).[3]

| | | |
|---|---|---|
| A/B/C | P33316 | DUT |
| | P62633 | CNBP |
| | Q05682 | CALD1 |
| | P48634 | PRRC2A |
| | O95817 | BAG3 |
| | P28340 | POLD1 |
| A/C | Q9NZB2 | FAM120A |
| | Q15149 | PLEC |
| | P43487 | RANBP1 |
| | Q99426 | TBCB |
| | P60709 | ACTB |
| B/C | O14979 | HNRNPDL |
| | Q09666 | AHNAK |
| | P02545 | LMNA |
| | Q96AE4 | FUBP1 |
| | Q15637 | SF1 |

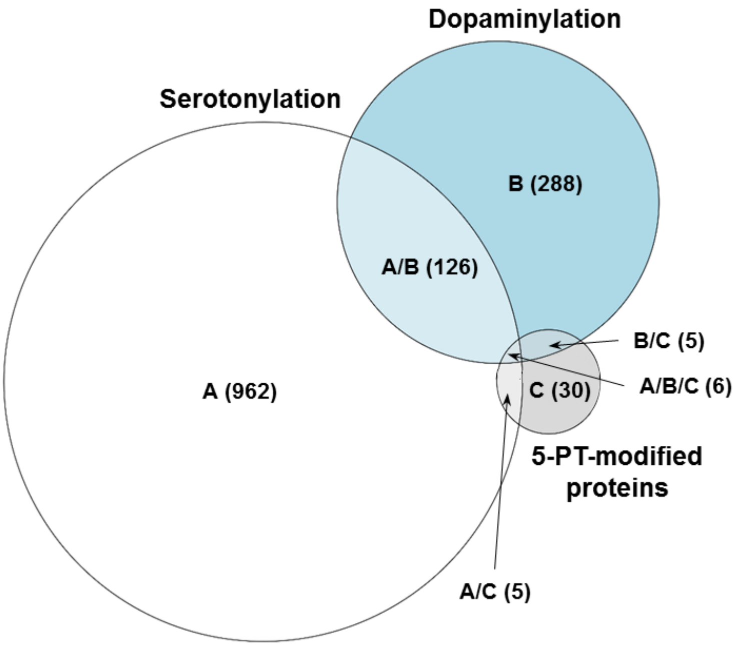

**Figure S6.** Uncropped immunoblotting images of Figure 4C, directly exported from the Odyssey CLx Imaging System (Li-Cor).

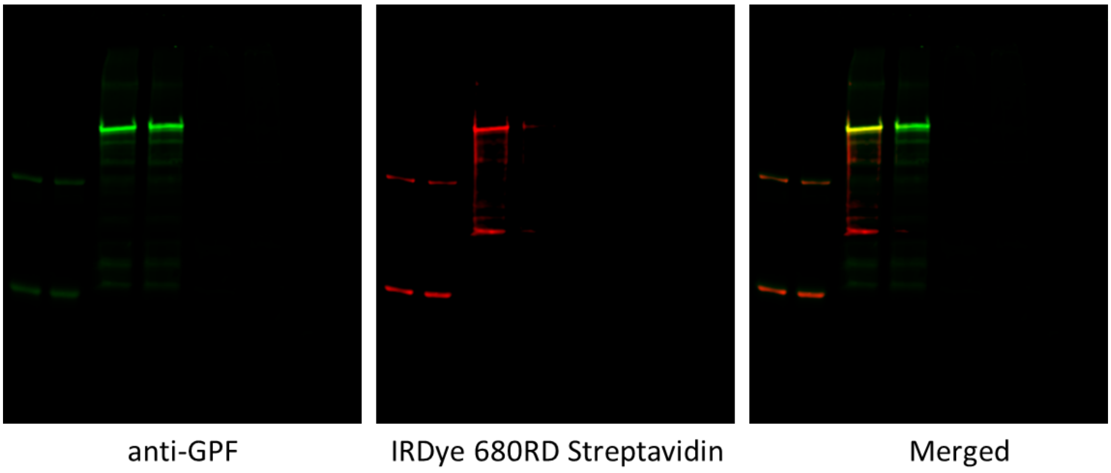

### REFERENCES AND NOTES

1. Zhang, N.; Wu, J.; Hossain, F.; Peng, H.; Li, H.; Gibson, C.; Chen, M.; Zhang, H.; Gao, S.; Zheng, X.; Wang, Y.; Zhu, J.; Wang, J. J.; Maze, I.; Zheng, Q. Bioorthogonal labeling and enrichment of histone monoaminylation reveal its accumulation and regulatory function in cancer cell chromatin. *bioRxiv* **2024**, doi: 10.1101/2024.03.20.586010.

2. Zhang, N.; Gao, S.; Peng, H.; Wu, J.; Li, H.; Gibson, C.; Wu, S.; Zhu, J.; Zheng, Q. Chemical proteomic profiling of protein dopaminylation in colorectal cancer cells. *bioRxiv* **2024**, doi: 10.1101/2024.04.27.591460.

3. Lin, J. C.; Chou, C. C.; Tu, Z.; Yeh, L. F.; Wu, S. C.; Khoo, K. H.; Lin C. H. Characterization of protein serotonylation via bioorthogonal labeling and enrichment. *J. Proteome. Res.* **2014**, *13*, 3523.
